# Personal genomics: new concepts for future community data banks

**DOI:** 10.1101/230516

**Authors:** Guy Dodin

## INTRODUCTION

The advent of fast, low-cost sequencing techniques is boosting the number of sequenced human genomes thus driving the irresistible push towards personalized genomics and personalized medicine.^1^ The examination of the genomes of large cohorts of patients will permit to identify the origins of many diseases and to propose adapted, fine-tuned medical treatments. The genomic boom is also expected to generate an economic activity worth billions of dollars and a fierce race has already started between Internet giants like Google and Amazon. Big data companies have already gathered tens of thousands of individual human genomes hosted in the cloud (that is on a server accessible via the Internet) and there processed by the bank’s customers (without sequence data transfer).This oligopolistic situation does not go without raising concerns when it comes to the cost and the quality of the services offered to the community of users (research institutes, hospitals, pharmaceutical companies, individuals) so that it would be advisable to open the field to a larger group of genomic service providers. Individuals, in growing numbers, are ready to consent financial efforts to have their own genome sequenced and, as a consequence, not only they will not accept additional expenses for storage of their data (as it is presently) but, they would even expect to be remunerated for handing out their personal details.

How safe and anonymous is information storage and how it can be controlled is another issue for individual s who accept to deposit their personal data in a gene bank. Scores of algorithms have been devised to ensure a satisfactory level of protection of both textual (name of persons, their attached medical files, etc..) and sequence data ^2,3^ and cases like the emblematic example of security failure illustrated by re-identification of anonymous donors through crossing freely available partial genomic information and family names are now readily circumvented.^4^ However, even encrypted,^7,8^ information will remain accessible to intruders due to the intrinsic limits of cipher protocols and to progress in computer technology.

In order to reconcile the security, control by their owners and availability to the scientific community of genomic data,^5,6^ and most importantly, to limit the possibilities of constitution of dominant oligopolies, a new type of genomic data bank is proposed here The project is organized along the following general lines. The bank is a community (mutual) bank (CB), that is, it is owned by those individuals depositing there the nucleotide sequences of their genomes. The CB is a repository site, whose role is limited to providing classical genomic banks, GenB (public or commercial) with the data their clients would subsequently handle and analyze *in situ*. The CB is organized along new lines in order to address security and access issues underlined above.

First, data are split into several parts independently stored. The information content of each piece is low enough so that breaking the confidentiality and the integrity of one will not allow the full information to be retrieved. In this context, data no longer need to be encrypted thus allowing easier handling. Importantly, split information permits to dispatch data to GenBs in a way that will promote competition for better service.

Second, the traffic of information and data between the CB and service providing gene banks, GenB, is managed and recorded in a blockchain.

## SPLIT GENOMIC DATA

A prior requirement is that access to one piece of split data (or several up to a certain limit) will never deliver enough information to unravel the genomic sequence stored in the CB. How genomic information can be made <u>surely incomplete</u>? A nucleotide sequence can be ‘vertically’ chopped into n subsequences each being subsequently separately stored. Although it assures confidentiality (access to one or several subsequences does not reveal the full information), this approach is unable to readily detect breaches of integrity stemming from addition, deletion, substitution of nucleotides. Alternatively, a sequence can also be split ‘horizontally’ that is, a sequence like ATTCAGC can be represented as 4 files (subsequently referred to as Base-files): ANNNANN, NTTNNNN, NNNNNGN, NNNCNNC. The literal Base-files can be replaced by binary sequences, called bin (Base) sequences, where, for example, the A-file leads to bin(A)=1000100, the T-file to bin(T) =0110000. This operation is not simply swapping symbols but rather it permits to transform letter sequences into numerical objects now amenable to processing by formal tools among them, Boolean algebra, linear algebra, Fourier Transform. Boolean OR and XOR (exclusive OR) operations on binary sequences, and binary array representations of a sequence readily allow detecting breaches in data integrity and easy reconstruction of the literal genomic sequence. The binaries can also be compressed as decimal or hexadecimal numbers, for more compact storage.

### Properties of the binary files

#### Confidentiality

Is a single binary sequence or the union of two enough to reconstruct the full genomic sequence or at least to provide sufficient information to break confidentiality? An illustration is provided by the following example. In a 100 base-long nucleotide sequence consisting of equal proportion of A, T, G, C, each binary sequence has about 25 ‘**1**’ and 75 ‘**0**’. The number of possible nucleotide sequences with the <u>same bin(Base) (</u>that is the number of colliding sequences) would be equal to 3^75^, and hence the probability on finding the true sequence by chance is (3^75^)^−1^ The union of two bin files, given their orthogonality (if there is **1** at a given position in any binary, the same position in the other files can only be occupied by **0**) results in a sequence with approximately 50 ‘**1**’ and 50 ‘**0**’. Hence, in this case the number of colliding sequences is 2^50^ and the probability of finding the genuine 100-base is 9.10^−16^. Supplying a third binary sequence makes this probability jump to 1 (1^25^) meaning full knowledge of the sequence. Those probabilities become vanishingly low with longer sequences (for a 1000-base sequence the above values will respectively be (3^750^)^−1^ and (2^500^)^−1^.

Even if its probability becomes significant (for short sequences), finding the true nucleotide arrangement by chance is unlikely because of the absence of a criteria to decide if the choice of one particular sequence among several others is the right one. If the ciphertext ‘Xc12VF’ is decrypted as ‘genome‘, speakers in numerous languages will immediately recognize this word as the true plaintext. Even if the ciphertext is only partially decrypted as ‘g-no-e’, an observer, familiar with the Latin alphabet of 26 letters, will make the reasonable choice, that among the 26^2^ possibilities, most of them leading to meaningless words, ‘genome’ is likely to be the true plaintext. Such a criteria for filling blanks does not exist for a partially decrypted sequence such as ‘AT-GC-’ since, in most cases, choosing ATTGCT, for instance, does not make more sense than selecting ATCGCT or any other sequence occurring with the same probability. Even a brute force approach where all blanks are replaced by any of the 4 nucleotides will not lift the indetermination. Hence, protection will still be assured for short genomic sequences. (Figure 1).

**Figure 1:**
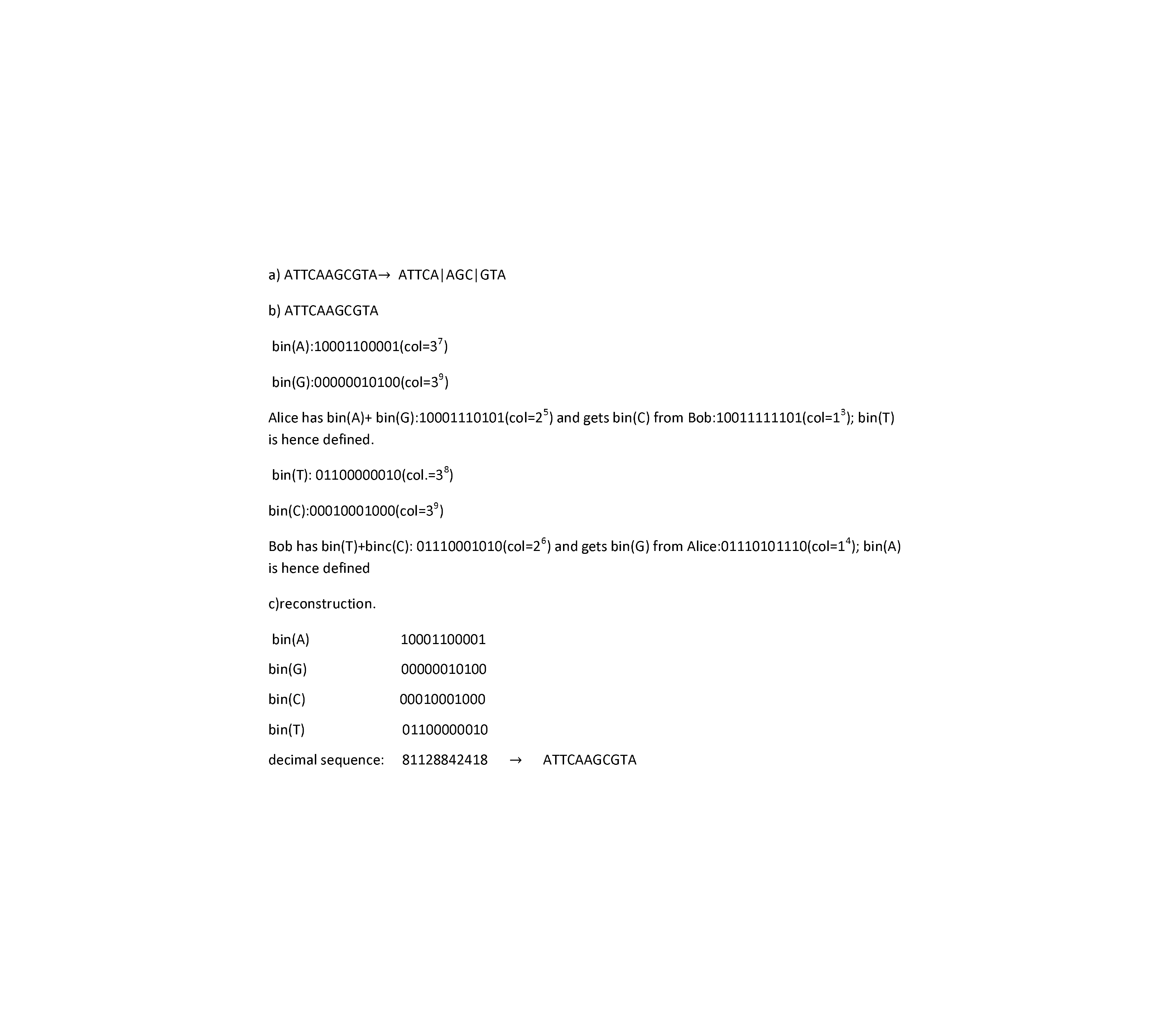
Partial information; a) vertical splitting; b) horizontal splitting; the number of collisions, col, is = (length of the residual alphabet)^number of 0^. c) sequence reconstruction from the binary rectangular array; the columns of the array represent decimal numbers

#### Integrity

When data are stored as literal sequences, corruption (replacement, deletion or addition of bases) remain undetected because there is no reference for what is the ‘true’ sequence. In contrast, the binary file representation, in many cases, gives an inner warning when corruption has occurred. Modification in the length of a binary file (insertion or deletion) is simply detected from comparison with the length of other binary files (supposedly not all corrupted).

Due to orthogonality, union of all binaries should give the complete file 111111111….When dealing with uncorrupted files, both Boolean term by term OR and XOR operations yield the same binary. If now a 0 is replaced by a 1 in a corrupted file, one of the binaries resulting from ORing must be different from that from XORing (one extra 0 appears in the XORed file, with respect to the ORed one). If a 0 replaces a 1, the overall union has one 0 (it should have none). (Figure 2)

**Figure 2:**
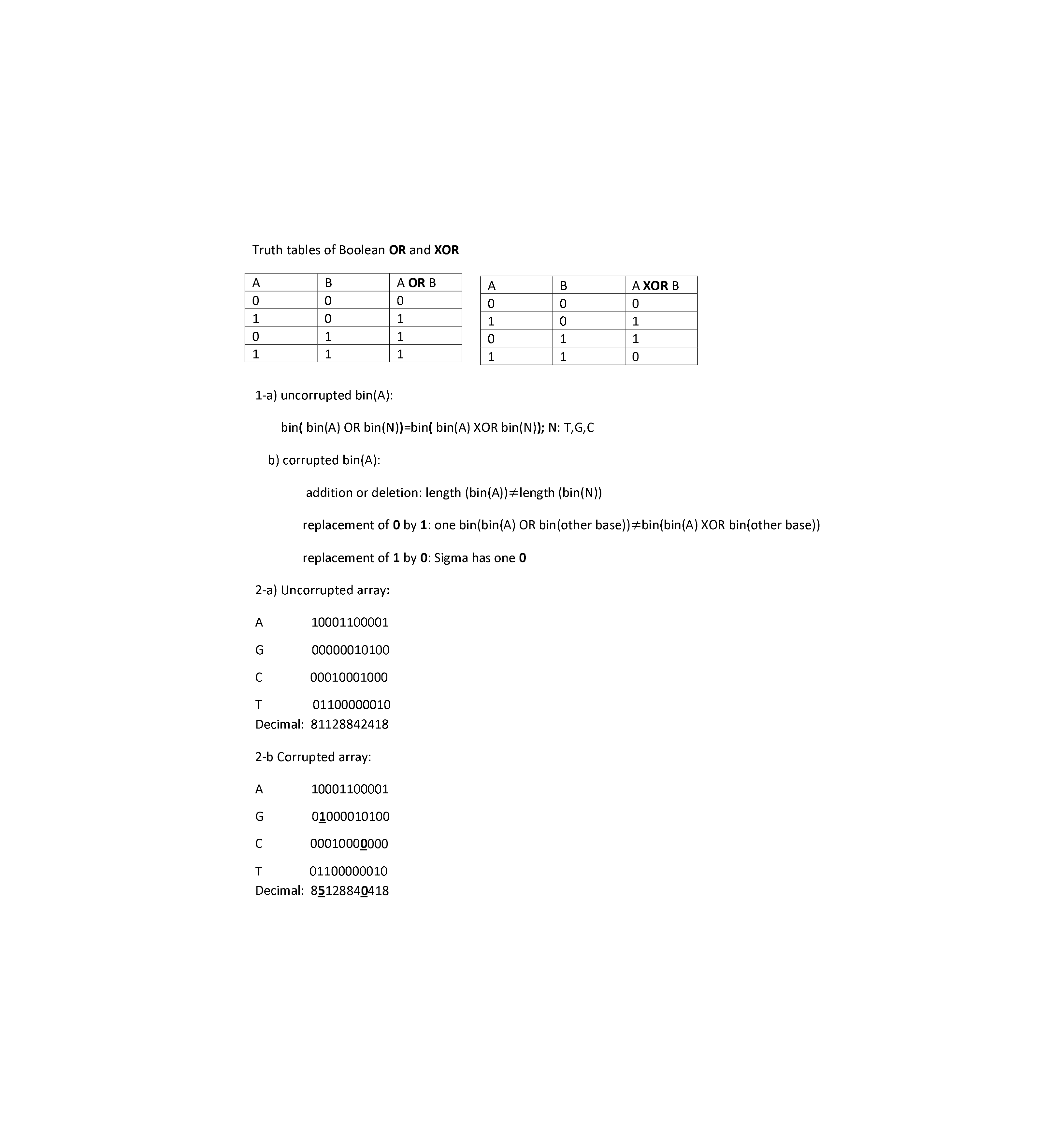
detection of failed integrity of binary files 1) illustrated with bin(A) and uncorrupted bin(G); Sigma is the union of all bin(Base) and is binary (11111111…..).;2) illustrated with the overall binary array

Moreover, any genomic sequence can be represented as a unique rectangular binary array of 4 lines whose lengths are that of the genomic sequence. Due to the orthogonality of the bin(Base) files, each column in the array represents one of the decimals, 1 (2^0^), 2 (2^1^), 4 (2^2^), 8 (2^3^). Thus, occurrence of any digit different from those in the decimal sequence indicates file corruption. In passing it is worth noting that the resulting unique decimal sequence (when uncorrupted) allows straightforward reconstruction of the nucleotide arrangement. (Figure 2)

Hence, failed integrity (even left undetected) has no consequence on confidentiality of the sequences when they are represented as binary files.

## TRANSACTIONS BETWEEN COMMUNITY (CB) AND SERVICE PROVIDERS BANKS (GenB)

Within the new project, if a GenB wants to gain full knowledge of a nucleotide sequence it will have to buy it from the community bank The general procedure, here illustrated with the simple example of 4 participating GenBs can be readily extended to any number of partners by sub-splitting the binary files, and is as follows.

As seen earlier, 3 binary sequences are necessary (and sufficient) to reconstruct the full genomic sequence. A GenB can acquire the 3 bin sequences but if it wants to access the genomes of a significantly large cohort of individuals it will be facing prohibitively high expenses especially if the community bank asks for a high price for a single binary. What the GenB can do is to exchange a copy of the bin(Base) it has bought with the copy of a complementary bin(Base) acquired by another GenB. Iterating the process over all partners, each GenB will get 3 bin(Base) (that is the full sequence) for the price of one. The community bank can even decide to refund (totally or partially) GenBs for their expenses if evidence for exchange deals concluded between partners is provided (for example if each GenB transmits back 3 bin(Base) sequences or parts of them, to the community bank). Hence, eventually, genomes can be accessed at <u>no cost</u> for those GenBs accepting to share their data.

The benefits mutual banks are expected to provide are guaranteed only if a large majority of individuals accepts to deposit there their personal data in order for CBs to eventually become the primary purveyors of genomic sequences. How would be organized a mutual bank to insure control, security and confidentiality genome depositors are expecting? The bank will have to manage the influx of data coming from sequencing facilities and their outwards transfer to clients GenBs along the split data protocol. A blockchain (BC) appears to be appropriate for controlling these exchanges. The core of a BC is the so-called ‘smart contract’, SC, where the conditions ruling asset transfers are hosted and automatically executed (without any external intervention). The results are recorded in a ledger together with other information like the time of the execution of the contract, the identification of the assets transferred (here genomic data), the financial balance of the bank. The ledger is permanently accessible to the members of the mutual bank and after being validated it is appended to previous blocks of information along a chain that represent the history of the bank activity.

## RESULTS AND DISCUSSION

### Data security

Security, that is confidentiality and integrity, of genomic data is satisfactorily insured by the bin(Base) representation of sequences. Is there any risk of unraveling the genuine sequence by alignment ^9^ with existing sequences stored in an open access library, by a GenB in the possession of only 1 bin(Base)? In an ideal situation, this should not occur since, as agreed by the community of depositors, gene sequences should not be found outside the new bank. Nevertheless, will security be at stake if data were hosted in open-access gene bank as a consequence of erroneous or malevolent operations? In order to illustrate such a situation the 4 Base-files, the literal equivalent of the bin(Base) file, were computed from human genomic sequences available in an open bank (sequences from chromosome 1 and chromosome Y burrowed from the ENSEMBL-BIOMART platform as examples). All sequence alignment algorithms employed failed to identify the original chromosome. Even the file resulting from addition of 2 Base-files did not permit identification. Although of no interest, the union of 3 Base-files successfully reveals the name of the chromosome.

Would a more in-depth inspection of a single or union of 2 bin(Base) files reveal insights that might allow identification of the genuine sequence?

Since they indicate the positions within the sequence of a particular nucleotide, the bin(Base) files capture part of the biological information conveyed by the nucleotide arrangement. For instance, from bin(G) and bin(C) one can localize CpG islands supposedly more frequent in gene-rich regions of human chromosomes. Also and according to the <u>second</u> Chargaff’s rule, the number of A and T (and G and C<u>) along a single strand</u> are roughly equivalent, and thus this might help inferring the overall number of ‘**0**’ or ‘**1**’ in the complementary binary sequence. Long range correlations observed in many genomes (including human), with additional short range triplet correlation in bacterial genomes, indicate periodic patterns in genomic sequences. In this respect, the Fourier spectrum of the bin(Base) files reveals, in the low-frequency domain, a long-range correlation ^10^ and, in the case of bacterial genomes an additional triplet correlation arising from intronless, densely packed protein-coding genes. The triplet correlation, revealed by the 1/3 frequency in the Fourier spectrum, is the signature of bacterial genomes which can be recognized as contaminants in complex, unidentified samples. All these periodical properties are too global to grant information at the single nucleotide level and hence they are not likely to undermine bin(Base) decomposition as a safe tool for confidentiality preservation.

### Community banks

Community (mutual) banks, yet to be installed, will act as buffers between individual raw genomic data just out of sequencing and classical gene banks, either public or private. Although data would also be hosted in the cloud (as separate binary files at different locations to preserve security) in contrast with general gene banks which trade with large numbers of users, the CBs will have only a limited number of customers (namely the GenBs themselves) and will not host facilities for their clients. As a consequence implementing a CB should only require limited computing resources for storage of data and management of the blockchain.

As amply stressed in this contribution, the motivations for promoting community banks are basically, the demand from genome depositors to exert control on their personal data (achieved by blockchain organization), and, most importantly, the necessity to avoid the genetic information to be concentrated within a small group of gene banks. A successful practical realization of the concept of CB requires people to join the project in great numbers.

